# SPARCS, a platform for genome-scale CRISPR screening for spatial cellular phenotypes

**DOI:** 10.1101/2023.06.01.542416

**Authors:** Niklas A. Schmacke, Sophia C. Mädler, Georg Wallmann, Andreas Metousis, Marleen Bérouti, Hartmann Harz, Heinrich Leonhardt, Matthias Mann, Veit Hornung

## Abstract

Forward genetic screening associates phenotypes with genotypes by randomly inducing mutations and then identifying those that result in phenotypic changes of interest. Here we present <u>spa</u>tially <u>r</u>esolved <u>C</u>RISPR <u>s</u>creening (SPARCS), a platform for microscopy-based genetic screening for spatial cellular phenotypes. SPARCS uses automated high-speed laser microdissection to physically isolate phenotypic variants *in situ* from virtually unlimited library sizes. We demonstrate the potential of SPARCS in a genome-wide CRISPR-KO screen on autophagosome formation in 40 million cells. Coupled to deep learning image analysis, SPARCS recovered almost all known macroautophagy genes in a single experiment and discovered a role for the ER-resident protein EI24 in autophagosome biogenesis. Harnessing the full power of advanced imaging technologies, SPARCS enables genome-wide forward genetic screening for diverse spatial phenotypes *in situ*.

## Introduction

Genetic screens offer a powerful approach to dissecting the complexity inherent in biological systems. Within this space, forward genetic screening is an unbiased way to map phenotypic changes to changes in the genome: From a library of genetic variants generated by random mutagenesis, mutants with interesting phenotypes are selected and their genotypes determined. This approach has led to groundbreaking discoveries in a variety of model organisms (*1–3*). Now, with the ability to specifically target mutagenesis to exonic regions of interest and disrupt both alleles of a given genetic locus, CRISPR-based genome editing technologies (*4*) have enabled the generation of large mutant libraries in which a single gene is knocked out in each cell (*5*). Individual genetically perturbed cells can now be profiled for their transcriptome (*6–10*), protein expression (*11*), spatial composition (*12*) and chromatin landscape (*13*). However, genome-wide screening libraries typically contain tens of millions of cells, a scale with which most of these techniques are currently incompatible. To overcome this limitation, only those cells with an interesting phenotype are typically isolated from the library and subsequently genotyped. This paradigm has largely limited cell-based genome-wide screens to three types of easily selectable phenotypes: a difference in proliferation rate, an inhibition of cell death, or a change in fluorescence intensity compatible with fluorescence-activated cell sorting (FACS) (*14–17*).

Increasingly powerful microscopic imaging provides information-rich data on diverse cellular phenotypes (*18*) and would therefore be an ideal technology to read out biological phenotypes of interest, particularly if combined with recent advances in deep learning. However, its application in genome-wide forward genetic screening has been hampered by a lack of scalability and other limitations: ‘in situ sequencing by synthesis’, a technology originally developed to profile the cellular transcriptome in tissue samples, has been adapted to sequencing short perturbation-encoding barcodes on the DNA level (*19, 20*). This method separates genotyping and image collection, resulting in complete image datasets for unbiased identification of new phenotypes. However, by design it does not include an enrichment step for selected phenotypes, requiring all cells in a mutant library to be sequenced irrespectively of whether they show a phenotype. In addition, the genotype can only be determined for a fraction of cells due to low sequencing fidelity even in low-complexity libraries (*20*), which in combination with the technology’s high costs has limited its applicability for screening genome-wide libraries at sufficient coverage (*21*). Image-based flow cytometers with sorting capabilities have recently enabled the investigation of spatial phenotypes at high throughput (*22, 23*). These devices rely on low-resolution flow-based microscopy of detached cells, preventing the identification of complex phenotypes. In addition, this technology makes sorting decisions in real time, restricting it to the identification of predefined phenotypes and preventing reanalysis of past screens. A method originally proposed for the transcriptomic characterization of B-cell populations (*24*) photoactivates fluorophores to mark cells for subsequent isolation by FACS (*25–27*). This approach can only separate few different phenotypes by fluorophore brightness (*28*). It also requires a real time decision on which cells to isolate, preventing whole-dataset analysis to discover unexpected new phenotypes and has not been demonstrated to be compatible with cell fixation, which is necessary for antibody-based staining of intracellular targets.

To enable robust genome-wide high-throughput screening for spatial cellular phenotypes, we set out to develop a technology that meets four key requirements: First, it should work on cells *in situ* and utilize state-of-the-art microscopy techniques. Second, it should accommodate large screening libraries to ensure adequate representation of rare phenotypes. Third, it should be compatible with the unbiased identification of previously unknown phenotypes from entire complex image datasets rather than single images in real time. Fourth, it should allow for reanalysis and reselection of cells for genotyping from previous archived screens. Importantly, the latter feature would allow the application of novel image analysis methods to previously performed screens as they become available.

## Results

### Spatial genotyping by laser microdissection

To analyze the spatial composition of tissues and clinical samples by mass spectrometry, we have been advancing workflows based on laser microdissection (LMD), a technique that uses a focused UV laser to cut out and collect arbitrary shapes from tissue sections (*29, 30*). In a most recent development, deep visual proteomics (DVP), we use LMD to excise defined tissue regions for subsequent proteomic characterization of individual cell types or extracellular zones by mass spectrometry (*31, 32*). We reasoned that the isolation of single phenotypically interesting cells from a pooled library by LMD would provide an ideal basis for a forward genetic screening technology for spatial phenotypes.

LMD requires samples to be present on a membrane that can be cut by a UV laser, so we first tested whether cells could be grown and imaged directly on such polymer membranes. Indeed, on polyphenylene sulfate (PPS) membranes, spinning disk confocal microscopy produced high-quality images that showed normal cellular morphology (fig. 1A). By segmenting these images into individual cells, we generated multi-channel perturbation image datasets from which we aimed to identify cells with phenotypes of interest for genotyping (fig. 1B). We then developed a rapid cutting protocol for LMD that is compatible with subsequent genotyping by minimizing autofocus time and optimizing the trade-off between laser speed and accuracy. Compressing the cutting path and leveraging the fact that it is sufficient to isolate nuclei to determine a cell’s CRISPR perturbation by sequencing, we ultimately reached a speed of 1,000 nuclei per hour.

**Figure 1.**
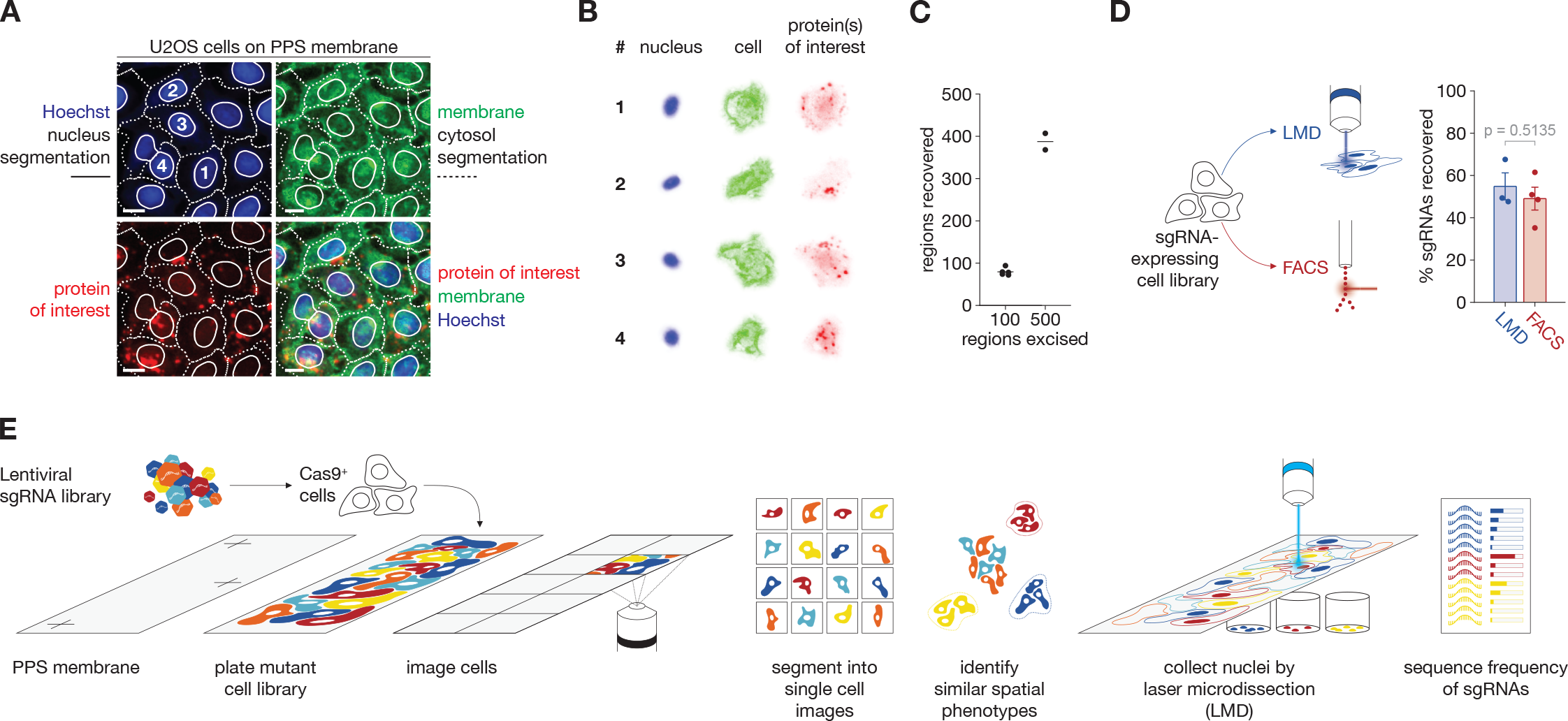
SPARCS enables genome-wide CRISPR screening for spatial phenotypes in human cells (A) Example images of U2OS cells on PPS membranes in several channels. Solid lines indicate nuclear segmentation based on Hoechst DNA staining; dotted lines indicate cytosol segmentation based on fast marching from nuclear centroids with wheat germ agglutinin (WGA)-Alexa 488 staining as a potential map. Numbers correspond to images of individual cells shown in (B). Images were acquired on an Opera Phenix microscope in confocal mode with 20 x magnification. Scalebars represent 15 µm. PPS: polyphenylene sulfide. (B) Post-segmentation images of individual mCherry-LC3 expressing U2OS cells. Numbers correspond to cells shown in (A). (C) 100 or 500 regions were excised from U2OS cells grown on a PPS membrane slide and subsequently counted. Five and two technical replicates were excised from one slide, respectively. (D) Comparison of sgRNA recovery after isolation of sgRNA-expressing fixed cells from one library either by laser microdissection (Leica LMD7, 1,000 nuclei per replicate, 3 independent biological replicates) or FACS (technical replicates). Bars indicate mean % sgRNAs recovered, error bars indicate SEM. p-value was calculated with an unpaired two-tailed t-test. (E) Overview of genome-wide CRISPR screening for microscopy-based spatial phenotypes with the SPARCS pipeline. Laser microdissection of individual nuclei on a Leica LMD7 has been optimized to isolate 1,000 nuclei/hr. Instructions for laser microdissection of selected cells are generated using our open-source python library py-lmd. PPS: polyphenylene sulfide.

Counting of excised membrane regions collected in a microwell plate using this protocol showed a yield of approximately 80 % (fig. 1C). We then tested the genotyping of excised nuclei by generating a pool of U2OS cells each expressing one of 77,441 unique sgRNAs in the Brunello CRISPR library (*33*), plated these cells onto PPS membranes and imaged them. To register membrane slides between imaging microscopes and the LMD microscope, we marked the membrane with calibration crosses as landmarks that define a coordinate system across each slide, allowing us to find the positions of cells to excise. We stitched individual field of view images of a slide into one whole slide image (WSI), segmented nuclei based on a DNA stain, generated a cutting map using our newly developed open-source python library py-lmd (fig. S1) and then excised and lysed 1,000 nuclei. Sequencing identified 549 unique sgRNAs on average in these lysates, demonstrating that isolating individual nuclei for subsequent CRISPR genotyping is feasible with LMD (fig. 1D). Comparing the number of unique sgRNAs in the LMD lysate with a lysate of cells from the same library isolated by FACS revealed that both techniques recovered an sgRNA from approximately 50 % of cells (fig. 1D). From these data we concluded that potential DNA damage induced by laser microdissection does not hamper sgRNA recovery. In summary, our results show that it is possible to employ LMD to recover genetic information from imaged cells at a throughput compatible with genetic screening (fig. 1E). We call this approach <u>spa</u>tially resolved <u>C</u>RISPR <u>s</u>creening (SPARCS).

### Validating SPARCS for genetic screening

To further develop SPARCS we applied it to screen for regulators of starvation-induced macroautophagy (hereafter referred to as autophagy), a fundamental process for cellular energy management (*34, 35*). The signature of autophagy is the formation of vesicles called autophagosomes. These are covalently decorated with proteins from the ATG8 family, including the well-studied human protein LC3B. During a key event in autophagosome biogenesis LC3B is conjugated to the head group of the lipid phosphatidylethanolamine (PE) through a series of ubiquitin ligation-like reactions. A critical component of this cascade is the protein ATG5 that forms an E3-like complex with ATG12 to mediate the covalent attachment of LC3B to PE. To follow the formation of autophagosomes during starvation we stably expressed LC3B tagged with mCherry in U2OS cells, because – unlike GFP – mCherry remains fluorescent upon fusion with the lysosome. We then treated these cells with the mTOR inhibitor Torin-1 to mimic starvation, which induces autophagy. Cells treated this way began to accumulate mCherry-LC3-positive puncta over the course of 14 hours (fig. 2A).

**Figure 2.**
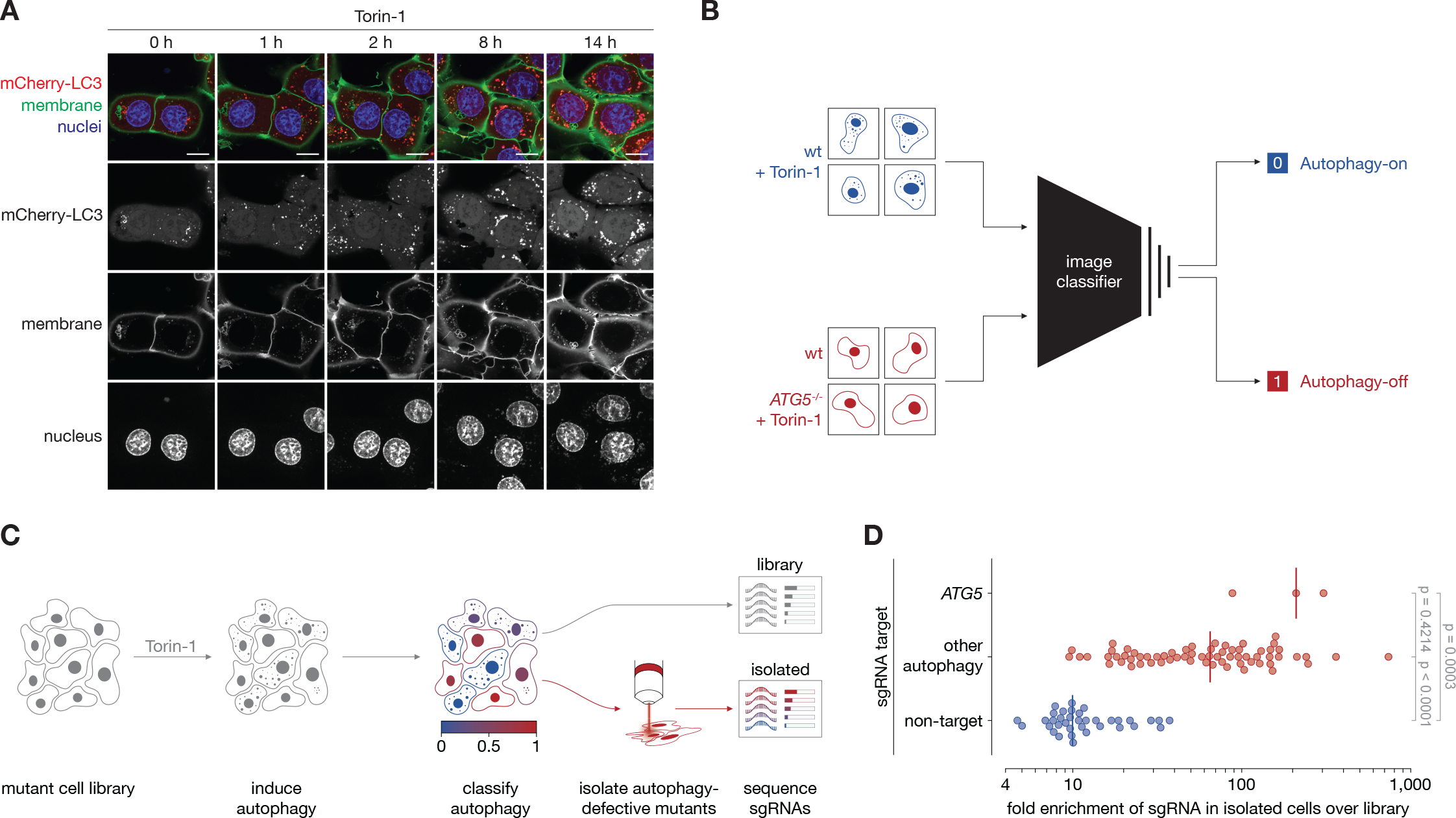
SPARCS achieves strong enrichment of spatial phenotypes in a forward genetic screen (A) U2OS cells expressing mCherry-LC3 and mNeon tagged with the lipidation signal of Lck at the N- terminus (membrane marker) were stimulated with Torin-1 and imaged live once per hour on a Nikon Eclipse Ti2 confocal microscope with 100 x magnification. Scalebars represent 15 µm. One representative of three independent experiments. (B) Schematic describing the training of a convolutional neural network-based image classifier for the identification of individual autophagy-defective cells. (C) Overview of SPARCS screening for autophagy. (D) Results from a SPARCS screen for autophagosome formation on 1.2 million U2OS cells. The top 0.1 % of cells classified as autophagy-off with a score above 0.94 by classifier 1.0 were isolated by laser microdissection (LMD) and their sgRNAs sequenced to determine their enrichment relative to the input library. p-values were calculated with a Kruskal-Wallis test followed by Tukey’s test.

In a screen, those cells containing sgRNAs against essential regulators of autophagy are unable to form these puncta. To identify these cells we trained a deep learning-based image classifier to differentiate between cells with or without autophagosomes (fig. 2B, fig. S2A). The training dataset was composed of segmented single cell images of mCherry-LC3 expressing U2OS cells that were treated with Torin- 1 (autophagy-on class) or left untreated (autophagy-off class). As an additional group we introduced cells treated with Torin-1, yet deficient in ATG5 (autophagy-off class). We used images from several biological replicates to avoid overfitting of our classifier to batch-specific characteristics such as staining intensity or cell density (table S1). To evaluate the performance of this classifier 1.0, we generated a new test dataset of images from unstimulated and Torin-1 stimulated wildtype and *ATG5*^-/-^ mCherry-LC3-expressing U2OS cells that had not been part of the training set and as such had never been seen by the classifier before. Classifier 1.0 achieved a false discovery rate (FDR) of < 1% (fig. S2B) at the chosen threshold, meaning that less than 1% of cells classified as potential hits with an autophagy-off phenotype were instead false positives that actually came from the autophagy-on class.

We then validated SPARCS by performing a small pilot screen on autophagosome formation in 1.2 million Torin-1-stimulated mCherry-LC3 U2OS cells transduced with the Brunello CRISPR knockout (KO) library (fig. 2C). From this library we isolated the top 0.1 % of cells classified as autophagy-off by classifier 1.0 with a score > 0.94, corresponding to a test set FDR of 0.38 %. Compared to the entire library, we found sgRNAs targeting *ATG5* to be highly enriched among isolated cells (median 200- fold) (fig. 2D). sgRNAs targeting other autophagy-related genes had a median of 60-fold enrichment with the most strongly enriched sgRNAs even exceeding 700-fold (fig. 2D). Control sgRNAs not targeting any human genes (‘non targeting controls’ (NTCs)) were rare among isolated cells with a median enrichment of 10-fold (fig. 2D). These results confirm that the SPARCS protocol stitches and registers WSI with sufficient accuracy for the isolation of the nuclei of interest. They also demonstrate that assessing autophagosome formation based on images is feasible with a deep learning classifier, and that in SPARCS, this classifier can be used to screen for autophagosome formation.

### Accurate detection of autophagy defects in single cell images

A classifiers’ Receiver Operating Characteristic (ROC) curve visualizes the tradeoff between true positive rate (the fraction of all autophagy-off cells that are correctly identified) and false positive rate (the fraction of autophagy-on cells incorrectly predicted as autophagy-off). The ROC curve of our classifier 1.0 confirmed its overall accuracy with an area under the curve (AUC) of > 0.92 (fig. S2C). However, at the precision (the fraction of predicted autophagy-off cells that are actually autophagy-off, 1-FDR) corresponding to 1 % FDR, the recall (= true positive rate) of classifier 1.0 was below 26% (fig. S2D). Closer analysis of the different categories of cells in the test dataset revealed that the classifier excelled at identifying autophagy-on cells, but performed poorly at recognizing autophagy- off cells (fig. S2E, F).

To improve classification of autophagy-off cells we refined our staining and imaging protocol and then trained a new version of our classifier. For this version 2.0 we decided to use a more streamlined multilayer perceptron (MLP) head with fewer trainable parameters, add another linear layer and increase the number of cells and biological replicates in the training dataset to capture as much biological variance as possible (fig. 3A, B, table S1). We also prefiltered the unstimulated and Torin-1 stimulated images for autophagy-off and -on cells to minimize the number of mislabeled training examples (table S1). To evaluate classifier 2.0, we first used parametric UMAP (*36*) to investigate if layers of the CNN had learned to differentiate between images of autophagy-on and autophagy-off cells.

**Figure 3.**
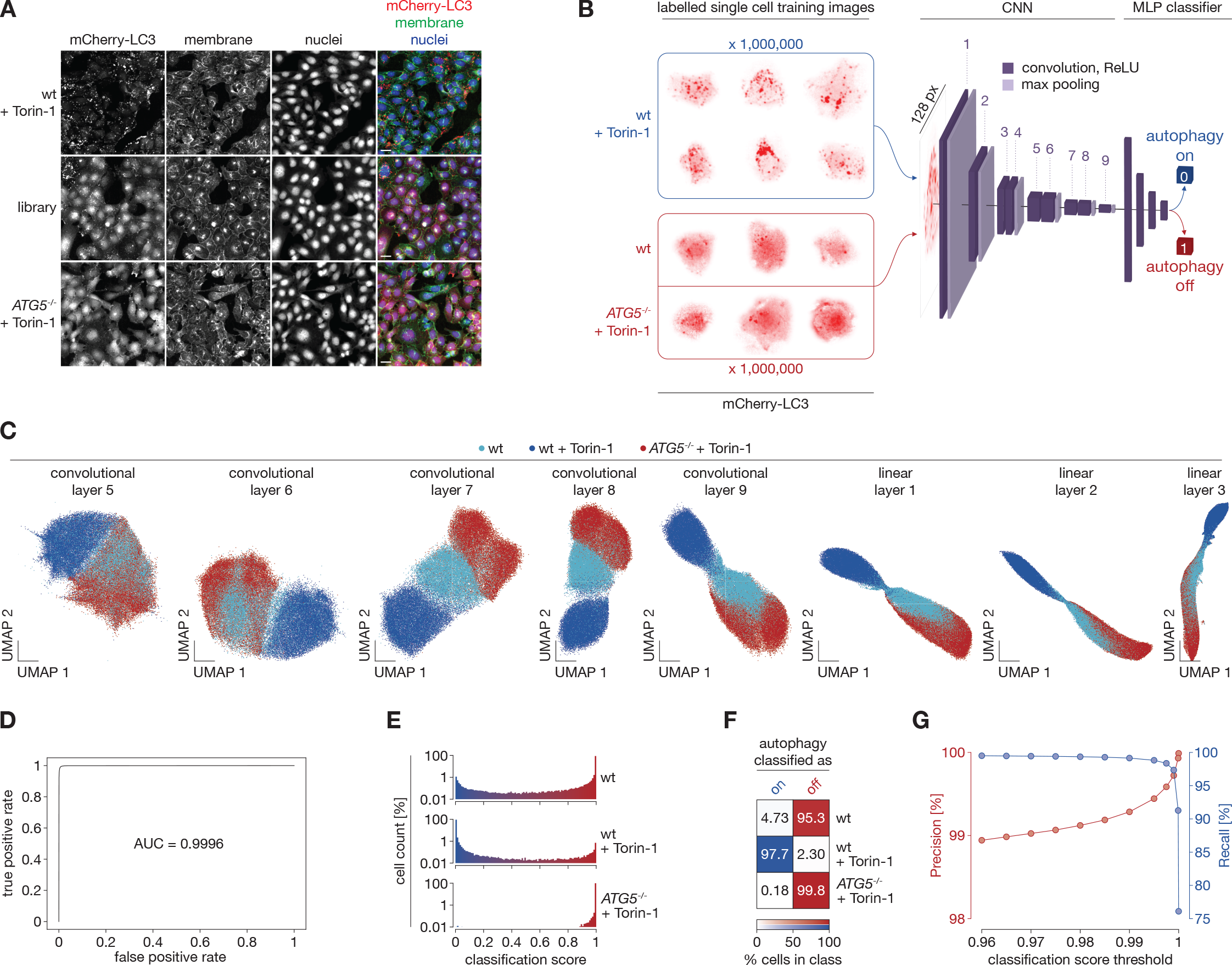
Deep learning accurately identifies autophagy-defective cells (A) Unsegmented images that were used for training autophagy classifier 2.0 after segmentation. Membranes were stained with WGA-Alexa488. “Library” refers to cells transduced with the Brunello CRISPR KO library. Images were acquired on an Opera Phenix microscope in confocal mode with 20 x magnification. Scale bars represent 30 µm. (B) Overview of the architecture and training paradigm of the convolutional neural network-based classifier 2.0 for autophagic or non-autophagic distribution of mCherry-LC3 in single U2OS cells. 1 million 128 × 128 px single cell images from several biological replicates were used in each training class. The autophagy-on class consisted of images of wildtype cells stimulated with Torin-1 pre-filtered for responsive cells. The autophagy-off class consisted of images of unstimulated wildtype cells pre- filtered to remove cells showing spontaneous autophagosome formation and images from two different *ATG5*^-/-^ clones. Images were acquired on an Opera Phenix microscope in confocal mode with 20 x magnification. CNN: convolutional neural network. MLP: multilayer perceptron. (C) UMAPs of mCherry-LC3 images of single U2OS cells featurized through the autophagy classifier 2.0 illustrated in (B) up to the indicated layers. Colors depict the indicated genotypes and treatments. 20,000 cells are shown for each genotype and treatment. (D) Receiver Operating Characteristic (ROC) curve of the autophagy classifier 2.0. AUC: Area under the curve. (E) Histograms of images of mCherry-LC3 expressing U2OS cells of the indicated genotypes treated as indicated after autophagy classification with our classifier 2.0 as illustrated in (B). (F) Heatmap showing the percentage of cells in e classified as autophagy-on or autophagy-off with a classification score threshold of 0.5. (G) Precision (Percent *ATG5*^-/-^ among cells classified as autophagy-off from an equal mix of Torin-1 stimulated wildtype cells and *ATG5*^-/-^ cells) and recall (Percent *ATG5*^-/-^ cells classified as autophagy- off) of our autophagy classifier at different thresholds for classifying cells as “autophagy-defective”. The data used for (C) – (G) come from an independent test dataset that was not used during training of the autophagy classifier.

This revealed that wildtype cells stimulated with Torin-1, unstimulated wildtype cells and *ATG5*^-/-^ cells clustered separately in representations of lower layers, most prominently in the 8^th^ of 9 convolutional layers (fig. 3C). These results suggested that our CNN had now learned to featurize images of LC3 distribution in a way that enables accurate classification of cells undergoing autophagy. Of note, the network of classifier 2.0 was capable of discriminating between *ATG5*^-/-^ and unstimulated wildtype cells despite those cells being in the same training class (fig. 3C), a clear improvement over classifier 1.0 (fig. S2G). Its ROC curve was also drastically improved with an AUC of > 0.999 (fig. 3D). Remarkably, in the binary classification output almost all cells were correctly classified according to their autophagy status even with a simple classification score threshold of 0.5 (fig. 3E, F). With classifier 2.0, classification thresholds > 0.98 produced FDRs of < 1 %, with higher thresholds reducing the FDR further without yet diminishing the excellent recall of nearly 100 % (fig. 3G, fig. S2H). Thus, for a complex biological process such as autophagy, training a CNN-based classifier on images from comparatively few biological replicates achieves excellent performance.

### Genome-wide autophagy screen with SPARCS

Encouraged by these results we used SPARCS to conduct a genome-wide screen on autophagosome formation. We screened a library of 40 million mCherry-LC3 expressing U2OS cells at a median coverage of 1,818 cells per gene in the human genome in batches of 5 and 35 million cells (fig. S3). Classifying autophagy based on the distribution of LC3 within the first batch showed that 0.56% of cells had a score > 0.98. We regarded these cells as potential autophagy-defective hits and, upon examining the 8^th^ CNN layer featurization of their LC3 distribution using parametric UMAP, found them to cluster separately from autophagy-on cells in the library with a classification score < 0.02 (fig. S4A, B). For genotyping we divided the hits into six bins according to their classification score (fig. S4C): The top bin represented a cutoff at which we found *ATG5*^-/-^ to be strongly enriched in our test dataset, whereas the second bin corresponded to unstimulated wildtype cells. Bins 3 – 6 contained the remaining potential hits with a roughly equal number of cells per bin. Zooming in on the 8^th^ CNN layer featurization of the LC3 distribution in the potential hits revealed that cells in bins 1 & 2 clustered separately from bins 3 - 6 (fig. S4D). This indicated that they contained different phenotypic variants with regard to their LC3 distribution, potentially corresponding to stronger defects in autophagosome formation. Indeed, we observed the fewest LC3 puncta in cells from bins 1 & 2 (fig. S4E).

For the second genome-wide screen batch we refined our classifier by more stringently selecting training examples of autophagy-on and -off cells, thereby further improving its overall performance (fig. S5). Using this new classifier 2.1 we obtained similar results from the second batch compared to 2.0 on the first screen batch: 1.40% of cells were classified as autophagy-off with a score above 0.98 and in their 8^th^ CNN layer featurization these cells again clustered separately from their autophagy-on counterparts with a score < 0.02 (fig. 4A, B). We therefore applied the same binning strategy to these images (fig. 4C), and, upon zooming in on their 8^th^ CNN layer featurization using parametric UMAP, found the separation of cells in bin 1 & 2 to be even more apparent than in the first batch (fig. 4D, E). We then isolated a total of 395,173 nuclei across both screen batches and sequenced their sgRNAs. Given their similarity on the phenotypic level we analyzed the genetic data of both batches together. All bins showed a marked enrichment of targeting over non-targeting sgRNAs and enrichment scores up to 600-fold, promising the identification of autophagy relevant genes (fig. 4F). In line with our FDR calculation (fig. 3G, fig. S5B) and our conclusions from the featurization of individual images (fig. 4D, fig. S4D), sgRNAs targeting genes known to be involved in autophagosome formation were most strongly enriched in bins 1 & 2 (fig. 4F). On the gene level *ATG5*, which our classifier was trained to identify, was among the most highly enriched genes in several bins, validating our supervised classification approach in the context of this large-scale screen (fig. 4G). The Brunello library targets each gene in the human genome with four sgRNAs. While in the higher score bins 1 & 2, genes with a high mean enrichment score had several sgRNAs enriched, in lower bins genes with a relatively high mean enrichment score often only had a single highly enriched sgRNA, indicating potential off-target effects. This prompted us to evaluate the number of sgRNAs enriched per gene as an alternative metric to score screening hits. Here we again found the strongest hits to contain mainly autophagy-related genes (fig. 4H). Taken together, these results establish that SPARCS is highly effective for large scale genetic screens on spatial phenotypes. Furthermore, despite the inherent complexity of image-based phenotypes, our supervised classifier facilitated the enrichment of a very small subset of individual cells with a genetically defined phenotype from a diverse genome-wide library of 40 million cells.

**Figure 4.**
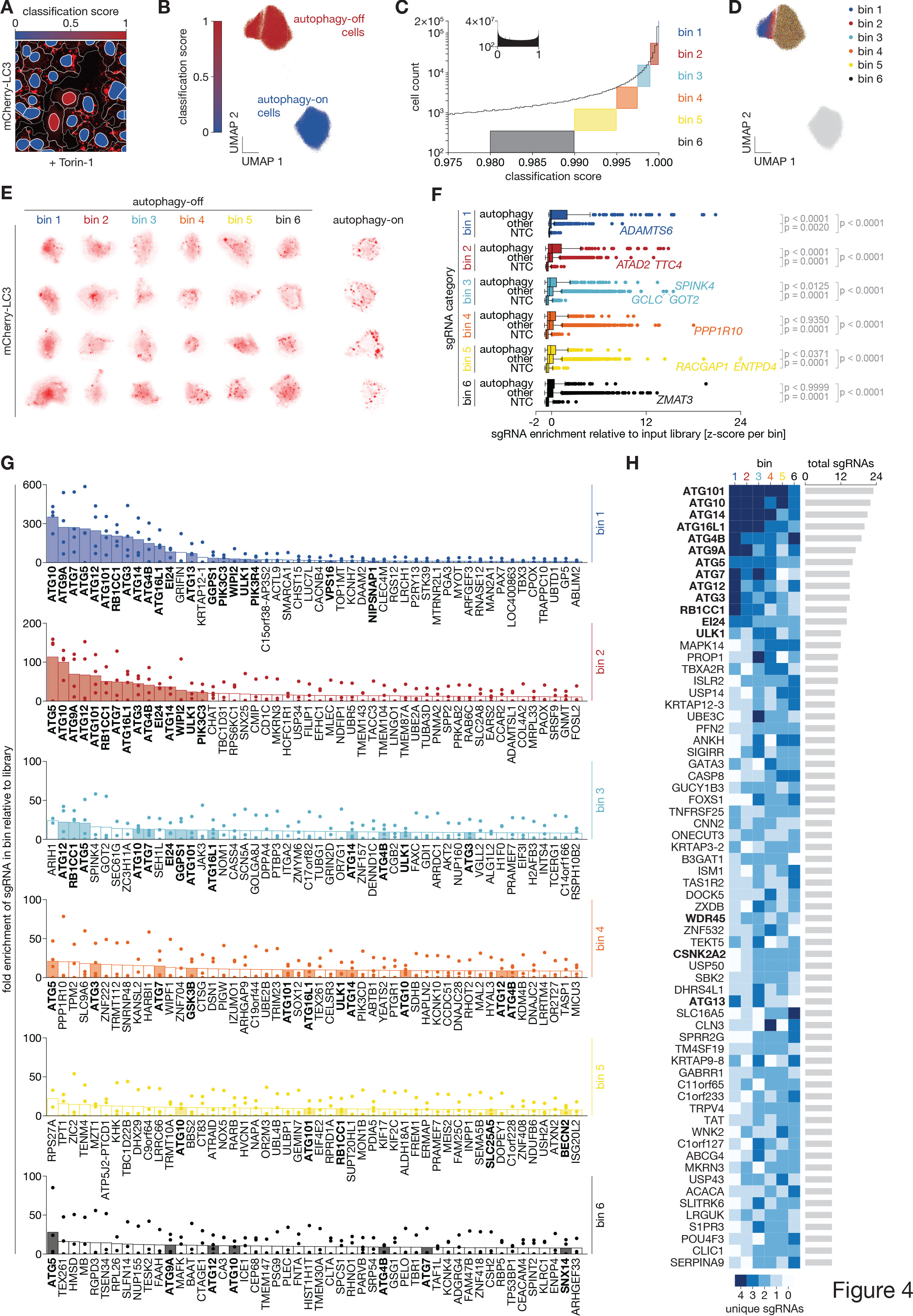
Genome-wide CRISPR screening for autophagosome formation in 40 million U2OS cells using SPARCS (A) Example region from a genome-wide SPARCS CRISPR knockout screen on autophagosome formation in mCherry-LC3 expressing U2OS cells after Torin-1 stimulation for 14 hrs. Colors in nuclei indicate the result of binary autophagy classification with the classifier 2.1, dotted lines indicate cytosol segmentation. Images were acquired on an Opera Phenix microscope in confocal mode with 20 x magnification. (B) Histogram of autophagy classification scores in the genome-wide CRISPR KO library batch 2 (inset) zoomed in on cells classified as autophagy-off with a score above 0.975. Colored boxes illustrate the binning strategy we used to isolate cells for sgRNA sequencing. (C) UMAP representation of single cell images from all cells in screen batch 2 with a classification score ≥ 0.98 (dark blue) or < 0.02 (light blue) featurized through the first 8 convolutional layers of autophagy classifier 2.1. 91,320 images are depicted for each category. (D) As C but colored according to our binning strategy along different autophagy classification thresholds as outlined in (B). 15,220 images are depicted per bin. Right panel shows a magnification of the UMAP region containing the putative screening hits. (E) Images of individual cells from each bin in screen batch 2. (F) z-scored enrichments of individual sgRNAs in each bin from batches 1 & 2. Vertical lines depict median, boxes depict interquartile range (IQR) and whiskers depict 1.5 × IQR. #: One sgRNA targeting the gene *ENTPD4* with a z-score of 42.1 in bin 5 is not depicted. p-values were calculated with a Kruskal-Wallis test followed by Dunn’s test. NTC: non-targeting control. (G) sgRNA sequencing results of the top 50 genes in each of the six bins filtered for genes for which we found at least two different sgRNAs in the respective bin in batches 1 & 2. Enrichment is calculated as the fraction of reads for an sgRNA in the respective bin divided by the fraction of reads of that sgRNA in the entire library. Bars indicate average enrichment per gene calculated from the enrichment of individual sgRNAs indicated as dots. Filled bars depict autophagy-related genes highlighted in bold. (H) Number of different sgRNAs per gene in each bin for all genes with at least 9 total sgRNAs across all bins. sgRNAs were counted if they were sequenced with a read fraction in the top 50 % per bin.

### EI24 reorganizes membranes for autophagy

The power of SPARCS became even more apparent when we evaluated our screen from the perspective of the investigated biological process: Remarkably, this single screen recovered almost all known essential genes of the starvation-induced macroautophagy pathway. This included the complete ULK1 complex and LC3 lipidation cascade (fig. 5A). Closer inspection of individual hits revealed that the most strongly enriched gene that is not a canonical macroautophagy gene was *EI24*, a gene coding for an ER-resident transmembrane protein (*37*) (fig. 5B). *EI24*^-/-^ cells have previously been described as autophagy-defective, but with a phenotype resulting in spontaneous LC3-puncta formation (*38*). This finding is not in line with the 82-fold enrichment of *EI24* in our screen, given that our classifier was trained to recognize cells with impaired rather than increased autophagosome formation. To investigate why we found EI24 KOs enriched among cells classified as autophagy-off, we generated individual *EI24*^-/-^ clones. Consistent with the previously reported spontaneous LC3 puncta formation, EI24- deficient cells have been described to exhibit increased lipidation of LC3 under steady state conditions (*38*), a phenotype we confirmed in *EI24*^-/-^ clones (fig. 5C). However, in contrast to previous results we found LC3 puncta formation in response to Torin-1 to be largely abolished in EI24-deficient cells (fig. 5D). Instead, these cells formed a single mCherry-LC3-positive speck that became more pronounced with Torin-1 stimulation, indicating a general defect in membrane traffic or autophagosome formation (fig. 5E). These results explain why our classifier picked up *EI24* knockouts and demonstrate again that even supervised image classification is capable of identifying previously undescribed phenotypes. Our results further indicate that *EI24* is required for autophagosome formation and has a function beyond its recently described LC3 puncta promoting role in maintaining Ca^2+^ homeostasis across the ER membrane (*39*) that remains to be investigated.

**Figure 5.**
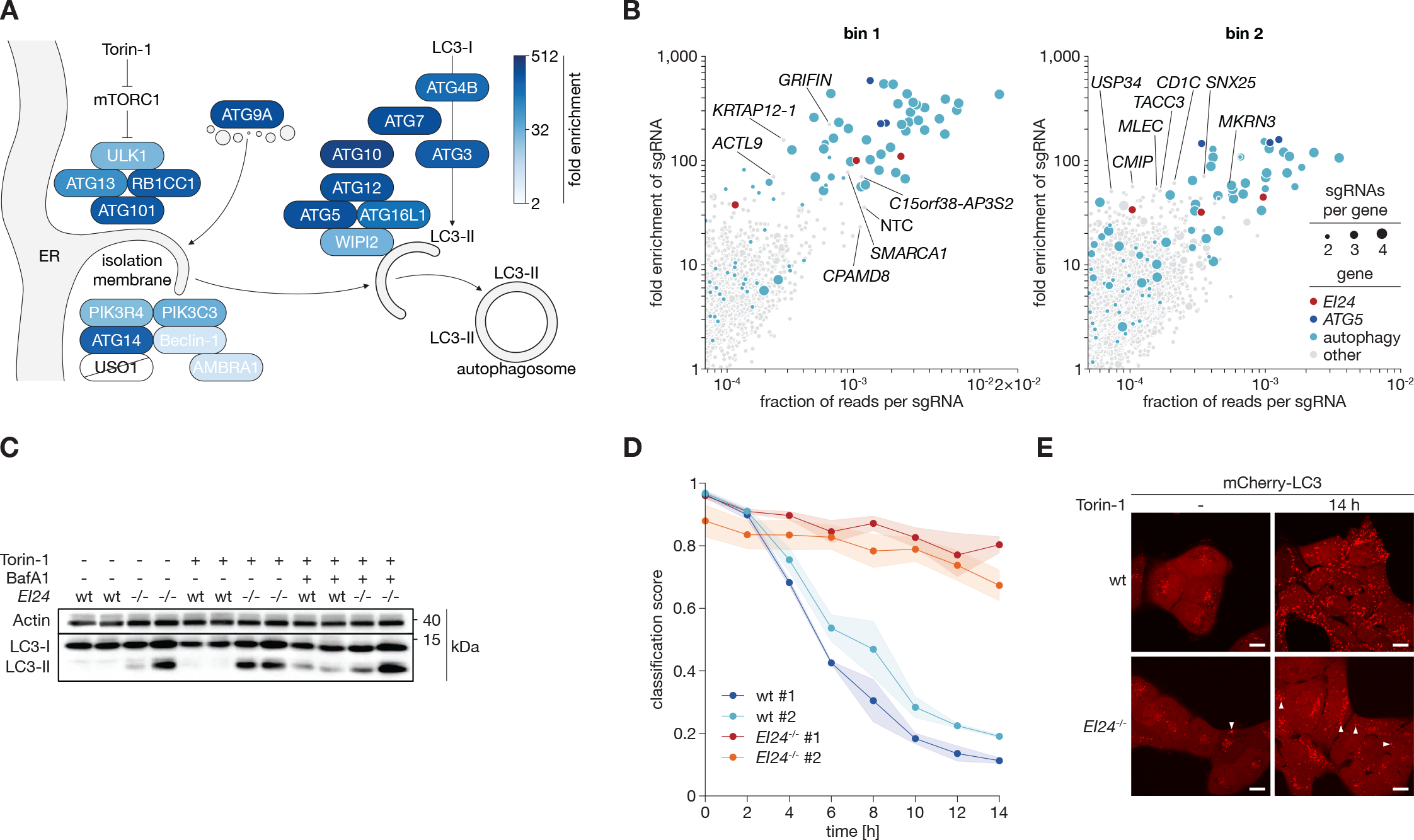
Analysis of hits from genome-wide SPARCS screen (A) Overview of the canonical macroautophagy pathway. Colors indicate the highest enrichment value we found for a given gene in any bin. *USO1* was not found with at least two different sgRNAs in any single bin. (B) Enrichment vs read count for individual sgRNAs in the top two bins for genes where we found at least two different sgRNAs in the respective bin. Circle sizes indicate the total number of different sgRNAs we found for a given gene, colors indicate different groups of genes. Individual sgRNAs from the “other” group are annotated. NTC: non-targeting control. (C) Immunoblot of endogenous LC3 lipidation in wildtype and *EI24*^-/-^ mCherry-LC3 and LckLip- mNeon expressing U2OS cell. Two clones are shown per genotype. One representative of three independent experiments. (D) Time course analysis of autophagy classification in clones of wildtype and *EI24*^-/-^ mCherry-LC3 and LckLip-mNeon expressing U2OS cells. Cells were treated with Torin-1 for up to 14 hrs. Dots represent average classifier scores from cells in 15 fields of view per timepoint and clone from three independent experiments, shaded areas represent SEM. Images were acquired on an Opera Phenix microscope in confocal mode with 20 x magnification. (E) Images of live mCherry-LC3 and LckLip-mNeon expressing wildtype and *EI24*^-/-^ U2OS cells after 14 hrs of Torin-1 stimulation. Arrowheads indicate larger mCherry-LC3 aggregates. Images were acquired on a Nikon Eclipse Ti2 confocal microscope with 100 x magnification. Scalebars represent 15µm. One representative of three independent experiments.

## Discussion

We present SPARCS, a platform that enables unbiased exploration of the genetic basis of subcellular spatial features in forward genetic screens. At the core of the SPARCS methodology, we have adapted and refined laser microdissection technology to unprecedented throughput to facilitate genetic screening applications. We have improved the precision and efficiency of isolating single nuclei from cell cultures, while automating the extraction of several hundred thousand nuclei into distinct bins. By integrating a deep learning-based classifier, our genome-wide SPARCS screen successfully identified nearly all known genes related to macroautophagy and revealed a novel phenotype associated with the *EI24* gene.

SPARCS offers a unique combination of features (table S2) that make it a powerful forward genetic screening platform. It can be seamlessly integrated with any state-of-the-art microscope for *in situ* cell imaging. The screening library size is not constrained, except by the imaging microscope’s throughput. Consequently, microscopy-based genome-wide perturbation screens can now achieve exceptional coverage. Besides the method described here, which involves isolating cells based on predefined classes, SPARCS is also compatible with the identification of individual cells exhibiting entirely novel or unexpected phenotypes. This is achieved through unbiased clustering and anomaly detection applied to the entire image dataset. Furthermore, we discovered that samples can be stored long-term, allowing for the reanalysis of archived SPARCS screens using newer algorithms. This facilitates the exploration of new biological insights within existing data. To streamline the process of translating the identification of individual cells with subcellular spatial phenotypes into a cutting map for LMD we have developed py-lmd, an open-source Python library for laser microdissection on arbitrary sample types that is available on GitHub. We hope that the accessible design of SPARCS, compatible with standard microscopes and sequencing workflows, will encourage its adoption by the scientific community.

Our screen uncovered a potential role in macroautophagy for *EI24*. This gene had previously been implicated in autophagy based on a *C. elegans* screen in 2010, but its mechanism of action had remained unclear (*38*). Beyond the original observation that EI24 deficiency leads to pronounced formation of non-functional autophagosomes even under steady state conditions, it was recently suggested that spontaneous Ca^2+^ fluxes across the ER membrane initiated autophagy in EI24 deficient cells (*39*). How spontaneous induction of autophagosome formation in *EI24*^-/-^ cells can be reconciled with a defect in autophagy remained unclear. The results from our screen and the following live cell imaging experiments, in which we found *EI24*^-/-^ cells to form fewer autophagosomes than wildtype cells, now suggest that EI24 plays a – potentially additional – role in autophagosome formation.

Systems biology is increasingly driven by large-scale artificial intelligence models that set new standards for reconstructing and predicting cellular behavior, but require enormous amounts of data to train. In light of this development, comprehensive, unbiased data acquisition approaches that can generate large datasets across modalities have become highly desirable. In this context, SPARCS, with its focus on open and accessible design and the ability to screen large libraries, can make a valuable contribution to understanding biology from the molecular to the organismic scale.

## Methods

### Cell culture

U2OS cells were cultured in DMEM supplemented with 10 % fetal calf serum (FCS), penicillin/streptomycin and 1 mM sodium pyruvate and split every 2-3 days. PPS membrane slides were sterilized for 30 minutes under the UV light of a cell culture hood with their cavity side down. Cells were then plated onto these slides cavity down in 4-well plates with 5 mL DMEM per well.

### Genome engineering

U2OS cells stably expressing Cas9, mCherry-LC3 and mNeon tagged N-terminally with the lipidation sequence of Lck (LckLip-mNeon, the original plasmid was a gift from Dorus Gadella (Addgene plasmid # 98821, (*40*))) were generated via lentiviral transduction. Briefly, HEK-293T cells were transfected with transfer plasmids for Cas9 or mCherry-LC3 and 3rd generation lentiviral particle production plasmids pMDLg and pRSV as wells as a VSV G-protein pseudotyping plasmid 18 hrs after plating. Eight hrs later, the medium was exchanged and cells were washed once in PBS. After 48 hrs supernatants containing viral particles were harvested and transferred onto U2OS cells plated 18 hrs before. 48 hrs later U2OS cells were washed. Cells were selected for Blasticidin resistance with 10 µg/mL Blasticidin or FACS-sorted for high fluorescent protein expression and single clones generated by limiting dilution cloning. A bright clone with a visible reaction to 600 nM Torin-1 was selected, expanded and used for all experiments. Lentiviral particles for the expression of individual sgRNAs from LentiGUIDE-Puro were generated analogously. Cells were selected for sgRNA expression with 5 µg/mL puromycin for 48 hrs. Of note, the cell line used for the autophagy screens did not yet stably express Cas9 but was instead transduced with a LentiCRISPRv2, a vector driving expression of both Cas9 and an sgRNA.

### Laser microdissection

Cutting paths for laser microdissection of selected cells were generated using our open-source python library py-lmd (https://github.com/MannLabs/py-lmd) with the configurations specified in the “screen config” file. Each shape was dilated to ensure that the cutting line did not go through or damage the nucleus. Laser microdissection was carried out on a Leica LMD7 at 40 x magnification using the software version 8.3.0.8275. The microscope was equipped with the Okolab LMD climate chamber (H201-ENCLOSURE-LMD and H201-LEICA-LMD) to ensure stable temperatures throughout the cutting process. Slides were equilibrated in the microscope to 34.5 °C before cutting to ensure focus stability. Cutting contours were imported from the XML files generated with py-lmd after reference point alignment and cut with the following settings: power 60, aperture 1, speed 100, head current 46 % - 51 %, pulse frequency 1128 and offset 185. Autofocus adjustment was performed every 30 shapes. Shapes were sorted into 48-well plates. During cutting a custom-built wind protection was used around the collection plate to ensure collection of excised shapes into the center of the well and prevent wind disturbances. After cutting, samples were stored at 4 °C before lysis and library generation.

### CNN-based image classifier training

Neural networks with 9 randomly initialized convolutional layers and 3 (classifiers 1, P) or 4 (classifiers 2.1 & 2.2) linear layers were trained to classify segmented single cell images as autophagy-on or autophagy-off (table S1). The training datasets were based on several biological replicates of mCherry- LC3 expressing U2OS cells with and without autophagosomes. The autophagy-off class consisted of images from unstimulated wildtype cells (pre-filtered to remove cells showing spontaneous autophagosome formation for 2.1 and 2.2) and two different *ATG5*^-/-^ clones. Where applicable, pre-filtering was performed with classifier P. The autophagy-on class consisted of single-cell images from stimulated wildtype cells, where applicable pre-filtered with classifier P to remove non-responding cells. To increase variability captured in the training data, the training slides were plated at an angle to include varying cell densities on one slide. 500 k, 1 million or 1.2 million single-cell images respectively were randomly selected from each class for training while ensuring balanced sampling from each test slide. An additional 50 k cells from each class were used for testing and validation during training. Training data were augmented by Gaussian blur, addition of Gaussian noise and random rotations in 90° steps. Training was performed using single-gradient descent with a learning rate of 1 × 10^-3^. Gradient clipping was set to 0.5. Training was performed over a total of 20, 30 or 40 epochs. Classifier performance was tested on a biologically independent set of unstimulated wildtype cells, Torin-1 stimulated wildtype cells and *ATG5*^-/-^ cells. Models were built and trained using PyTorch (*41*).

### Segmentation of individual cells

Images were flat-field corrected during image acquisition using the Perkin Elmer Harmony software (v4.9) and intensity rescaled to the 1 % and 99 % quantile. Extremely bright regions (pixel values greater than 40000) were set to 0 before determining the normalization quantiles. Stitching of image tiles was performed using the ashlar python API (*42*) in our open-source python library SPARCStools (https://github.com/MannLabs/SPARCStools).

Stitched whole slide images were segmented using our open-source SPARCSpy python library (https://github.com/MannLabs/SPARCSpy) with the parameters defined in “config_screen” or “config_training” respectively. A nucleus segmentation mask was generated using a local median thresholding approach and the cytosol segmentation mask was calculated using fast marching from nuclear centroids with WGA staining as a potential map.

Single cell images were extracted based on nuclear and cytosolic segmentation masks. The masked area was extended using a Gaussian filter with a sigma of 5 to extract information from each of the imaged channels and saved to hdf5 files as individual 128 × 128 px images.

### Sample preparation and imaging of autophagy

After stimulation with 600nM Torin-1, PPS slides were washed 1x in PBS in a Coplin jar and then stained with 10 µg/mL WGA-Alexa488 in PBS for 10 minutes at 37 °C. After washing 1 x with PBS slides were fixed for 10 minutes at room temperature in 4 % MeOH-free PFA in PBS in 4-well plates. After washing 3 x in PBS, slides were stained with 10 µg/mL Hoechst-33342 in PBS at 37 °C for at least 30 minutes. After washing 3 x in PBS, slides were dried in a centrifuge at 3,400 g for 1 minute. Cells in ibidi microwell slides and plates were stained according to the same protocol but imaged in PBS. Imaging was done on a Nikon Eclipse Ti2 spinning disk confocal microscope or an Opera Phenix high-content imager as indicated.

### Genetic screening for autophagy regulators

We conducted our screen in mCherry-LC3 expressing U2OS cells using the Brunello human CRISPR KO library in the LentiCRISPRv2 backbone. The Brunello library was a gift from David Root and John Doench (Addgene #73178) and amplified according to their protocol (*33*). U2OS cells were transduced with lentiviral particles produced as described above at an MOI of approximately 0.2. After 48 hrs, successfully transduced cells were selected with 5 µg/mL puromycin for two days and then expanded for three days. We then plated 50 million cells on a total of 109 slides in 4 well plates, and in addition included unstimulated and wildtype controls on separate slides with every screening batch for classifier training. The day after plating, cells were stimulated with 600 nM Torin-1 for 14 hrs. Slides were then prepared for microscopy as described above and imaged on an Opera Phenix high content imager at 20 x resolution. Where applicable slides were stored at -20 °C and brought to 4 °C the day before laser microdissection. Cells from each bin were excised into multiple wells. Nuclei were then lysed in 48- well plates using the arcturis PicoPure^TM^ DNA extraction kit. 120 µL of lysis buffer was added to each well and incubated at 65 °C for 4 hrs. Proteinase K was inactivated at 95 °C for 15 mins. Cooled samples were transferred to PCR tubes and the emptied wells were rinsed with 40 µL of ddH2O. Amplification of sgRNAs was performed as described previously (*43*) but in a single step PCR (*33*) over 36 cycles with no added water. Sequencing was performed on an Illumina NextSeq with 500 reads per nucleus on average. An sgRNA read count table was generated for each sequencing library. Low quality sgRNAs were removed by applying a minimum number of reads per sgRNA threshold that was set based on the distribution of read counts per sgRNA in the sample. The non-targeting sgRNA with the sequence TACGTCATTAAGAGTTCAAC was excluded from sequencing results of approximately 40 % of cells from bins 3 - 6 of batch 2 of the genome-wide screen due to a contamination of the sequencing library leading to abnormally high read counts of this specific sequence. For further analysis only sgRNAs with at least a fraction of reads corresponding to a single cell per well were used. sgRNA read fractions of individual wells were then aggregated per bin by multiplying with the fraction of excised cells in that well over all excised cells in the bin. The per-bin aggregated sequencing results were used for all further data analysis. Enrichment values were determined by normalizing the aggregated fraction of reads per sgRNA to the fraction of reads per sgRNA in the input cell library.

### Immunoblotting

20,000 U2OS cells were plated per 96-well. 18 hrs after plating, cells were stimulated and then harvested in 1 x Lämmli buffer. 3 wells were pooled per condition. Lysates were boiled at 95 °C for 5 min. and then run on 16 % TRIS-glycine polyacrylamide gels before immunoblotting onto 0.2 µm nitrocellulose membrane for 90 minutes. Membranes were blocked in 5 % milk in PBST for 1 hr and incubated with primary antibody at 4 °C overnight. After washing 3 x in PBST for a total of 15 min. membranes were incubated with HRP-labelled secondary antibody for 2 hrs at room temperature. After washing 3 x in PBST for a total of 15 min. membranes were covered in luminescent HRP substrate and immediately imaged.

## Acknowledgements

We would like to thank Larissa Hansbauer for outstanding technical support; Jochen Rech and the BioSysM Liquid Handling Unit for excellent support with robotics; Claudia Ludwig and the BioSysM FACS Core Facility for great support with cell sorting; Mario Oroshi and the computing centre of the Max Planck Institute of Biochemistry for computational support and IT infrastructure; the Imaging Facility of the MPI of Biochemistry and the Center for Advanced Light Microscopy (CALM) for support with light microscopy; Rin Ho Kim and the Sequencing Facility of the MPI of Biochemistry as well as Stefan Krebs and the Genomics unit of the Laboratory for Functional Genome Analysis (LAFUGA) for sequencing; and Falk Schlaudraff, Christoph Greb and Florian Hoffmann from Leica Microsystems for technical support.

## Author Contributions

Conceptualization: N.A.S., S.C.M.; Formal Analysis: N.A.S, S.C.M., G.W.; Funding Acquisition: M.M., V.H.; Investigation: N.A.S., S.C.M., G.W., A.M., M.B., H.H.; Resources: H.H., H.L., M.M., V.H.; Software: N.A.S., S.C.M., G.W.; Visualization: N.A.S., S.C.M., G.W.; Writing – original draft:N.A.S., S.C.M., M.M., V.H.; Writing – review & editing: N.A.S., S.C.M., G.W., A.M., M.B., H.H., H.L., M.M., V.H.

## Data and materials availability

Code to recreate the figure manuscripts is available on GitHub (https://github.com/MannLabs/SPARCS_pub_figures).

The code described in this manuscript is available from the following GitHub repositories:

The py-lmd python library provides code to direct excision of defined regions on a Leica LMD7 laser microdissection microscope (https://github.com/MannLabs/py-lmd).

The SPARCStools python library provides code to rename TIF image files generated by the PerkinElmer Harmony software and stitch these into WSIs (https://github.com/MannLabs/SPARCStools).

The SPARCSpy python library contains the autophagy classifiers, as well as code to segment and extract single-cell images from entire fields of view up to WSIs (https://github.com/MannLabs/SPARCSpy).

All other data are available from the authors upon request.

---

**Figure S1.**
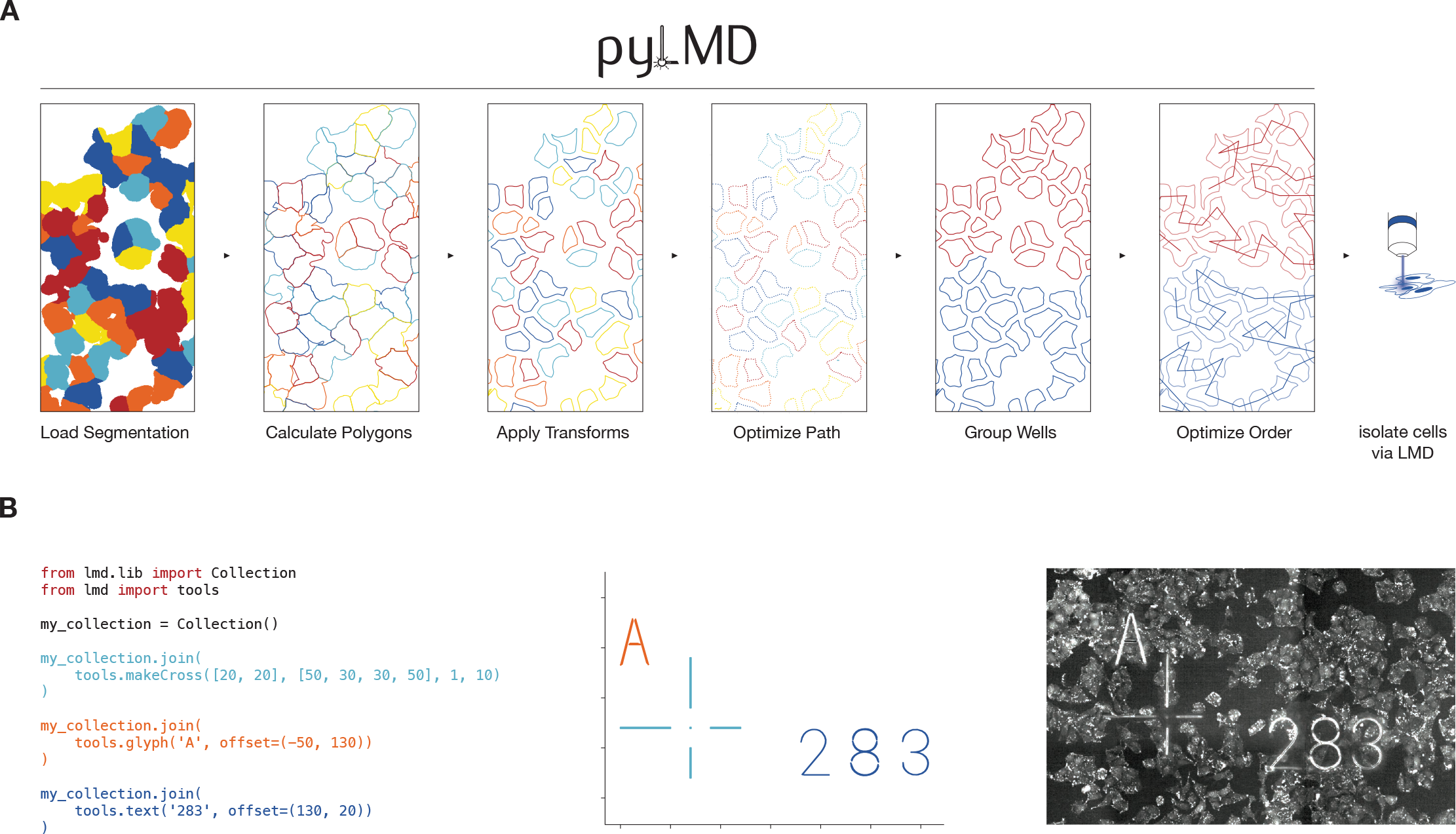
The py-lmd python library generates cutting paths for automated laser microdissection (A) Overview of cutting path generation with py-lmd. (B) py-lmd allows the generation of arbitrary shapes such as calibration crosses.

**Figure S2.**
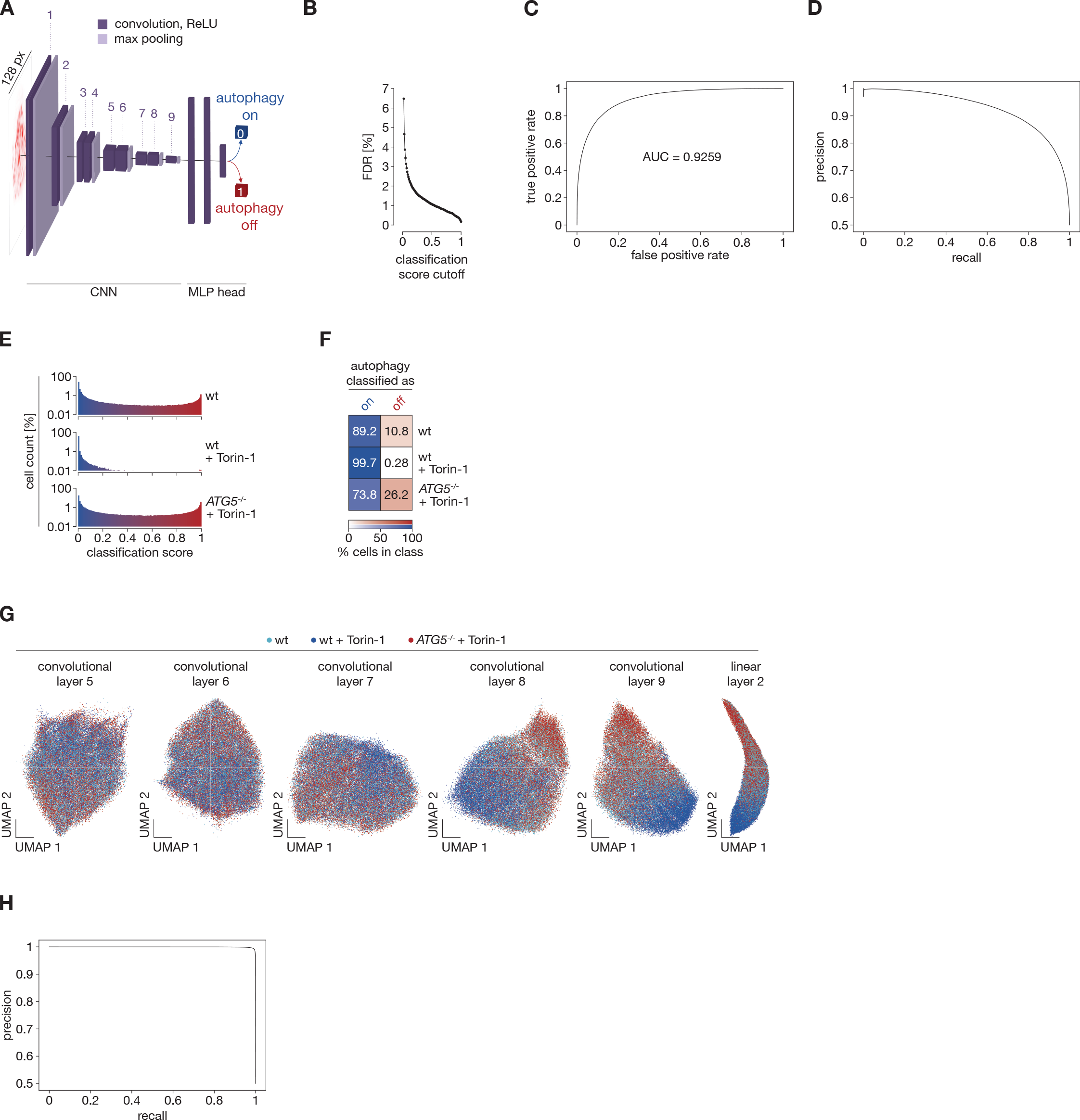
Performance of LC3 image-based autophagy classifiers (A) Overview of the architecture and training paradigm of our convolutional neural network-based classifier 1.0 for autophagic or non-autophagic distribution of mCherry-LC3 in single U2OS cells. 500,000 128 × 128 px single cell images from several biological replicates were used in each training class. The autophagy-on class consisted of images of wildtype cells stimulated with Torin-1. The autophagy-off class consisted of images of unstimulated wildtype cells and images from two different *ATG5*^-/-^ clones. CNN: convolutional neural network. MLP: multilayer perceptron. (B) False discovery rates (FDR) of the autophagy classifier 1.0 at different classification score cutoffs. (C) Receiver Operating Characteristic (ROC) curve for autophagy classifier 1.0. AUC: area under the curve. (D) Precision-Recall curve for our autophagy classifier 1.0. (E) Histograms of images of mCherry-LC3 expressing U2OS cells of the indicated genotypes treated as indicated after autophagy classification with classifier 1.0 as illustrated in (A). (F) Heatmap showing the percentage of cells in d classified as autophagy-on or autophagy-off with a classification score threshold of 0.5. (G) Parametric UMAPs of mCherry-LC3 images of single U2OS cells featurized through our autophagy classifier 1.0 illustrated in (A) up to the indicated layers. Colors depict the indicated genotypes and treatments. 20,000 cells are shown for each genotype and treatment. (H) Precision-Recall curve of classifier 2.0. (B) – (H) were calculated on independent test datasets for the respective classifiers

**Figure S3.**
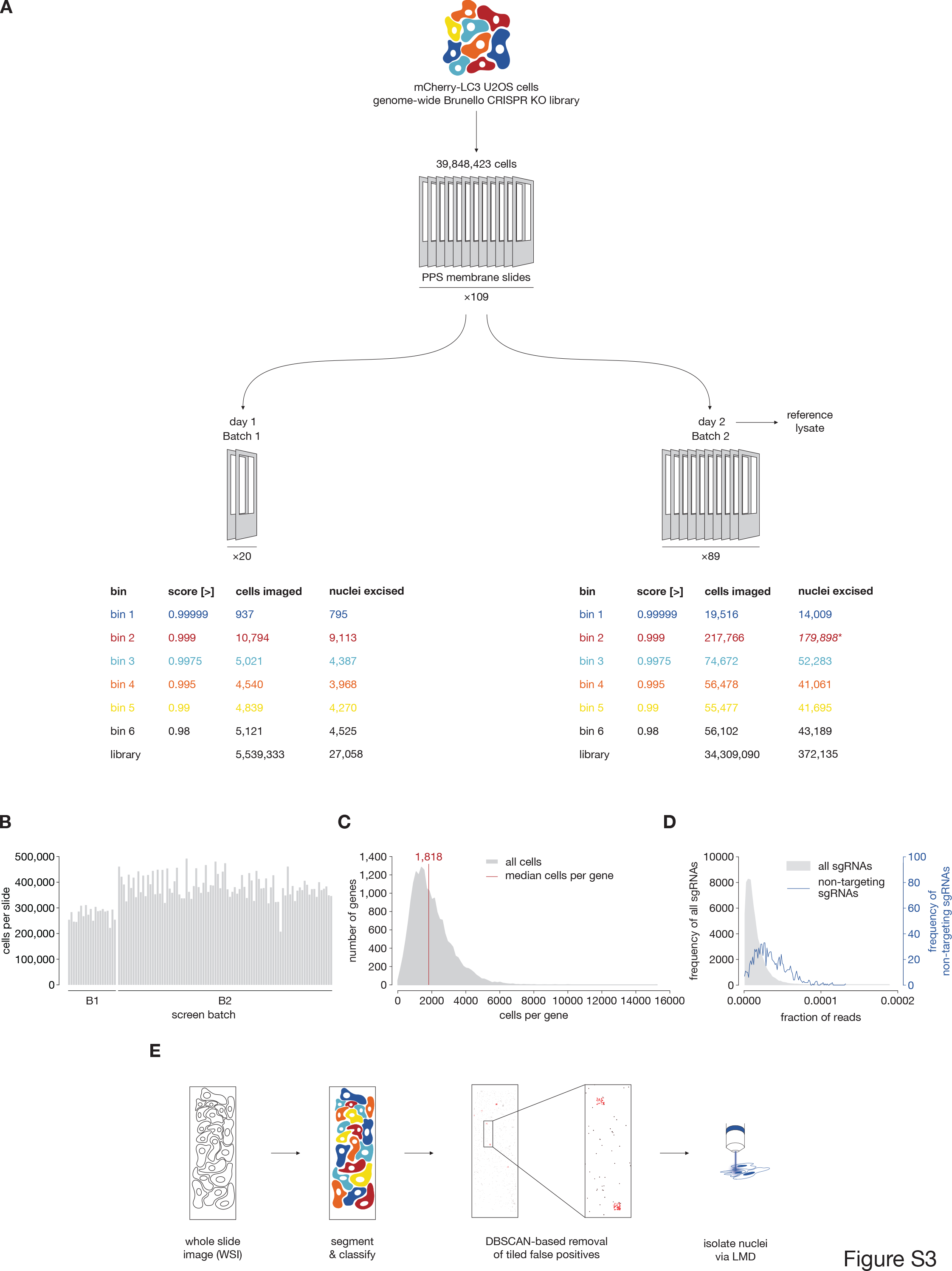
Overview of genome-wide SPARCS screening for autophagy (A) Batching and binning strategy for screening autophagosome formation in 40 million mCherry-LC3 expressing U2OS cells. We dissected fewer cells than we imaged for a given bin due to the quality control step outlined in e. *The efficiency of the PCR on bin 2 from batch 2 had decreased dramatically, presumably due to the high density of membrane fragments in the reaction, leading to a loss of sgRNAs for sequencing. PPS: polyphenylene sulfide. (B) Number of cells segmented per screen slide. (C) Distribution of human genes targeted in the screen across cells in the library as determined by deep sequencing. (D) Distribution of non-targeting and targeting sgRNAs in the reference library as determined by deep sequencing. (E) Quality control strategy for false positives arising from out-of-focus images. When we spatially clustered hits using DBSCAN, we found clusters above a certain size to correspond to entire out-of-focus imaging tiles and removed these clusters before nuclei excision.

**Figure S4.**
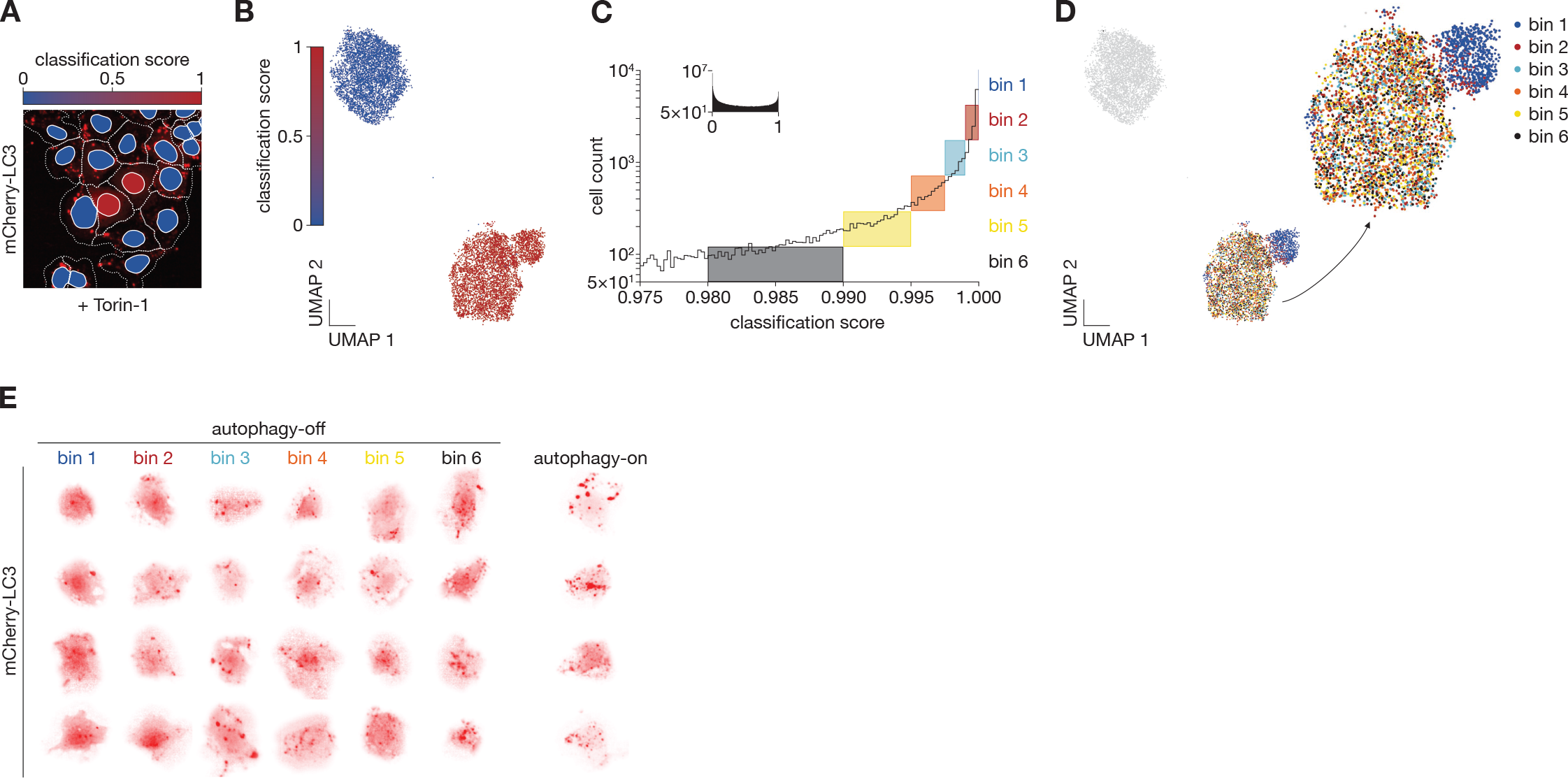
Results from genome-wide autophagy screen batch 1 (A) Example region from a genome-wide SPARCS CRISPR knockout screen on autophagosome formation in mCherry-LC3 expressing U2OS cells after Torin-1 stimulation for 14 hrs. Colors in nuclei indicate the result of binary autophagy classification with classifier 2.0, dotted lines indicate cytosol segmentation. Images were acquired on an Opera Phenix microscope in confocal mode with 20 x magnification. (B) UMAP representation of single cell images from all cells in screen batch 1 with a classification score ≥ 0.98 (dark blue) or < 0.02 (light blue) featurized through the first 8 convolutional layers of autophagy classifier 2.0. 4,806 cells are depicted per category. (C) Histogram of autophagy classification scores in the genome-wide CRISPR KO library batch 1 (inset) zoomed in on cells classified as autophagy-off with a score above 0.975. Colored boxes illustrate the binning strategy we used to isolate cells for sgRNA sequencing. (D) As (B) but colored by screening bin. 801 cells shown per bin. (E) Images of individual cells from each bin in screen batch 1.

**Figure S5.**
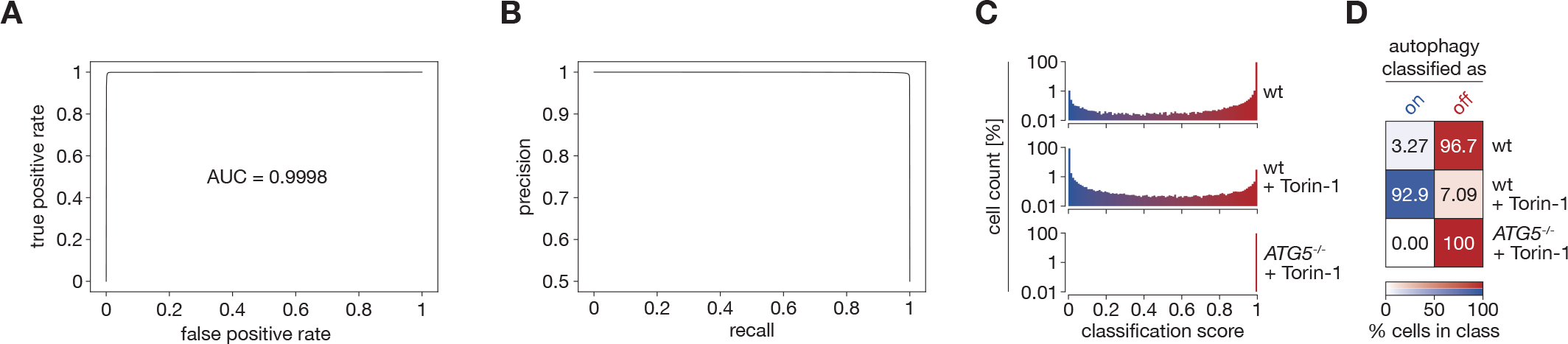
Performance metrics of classifier 2.1 (A) Receiver Operating Characteristic (ROC) curve for autophagy classifier 2.1. AUC: area under the curve. (B) Precision-Recall curve for our autophagy classifier 2.1. (C) Histograms of images of mCherry-LC3 expressing U2OS cells of the indicated genotypes treated as indicated after autophagy classification with classifier 2.1. (D) Heatmap showing the percentage of cells in (C) classified as autophagy-on or autophagy-off with a classification score threshold of 0.5. (A) – (D) were calculated on an independent test dataset.

**Table S1.** Overview of CNN-based image classifiers trained in this study All classifiers were trained on 128 × 128 px single cell images using PyTorch lightning. Unstimulated control slides containing library cells plated in parallel with the screen slides were included during training to capture possible batch effects introduced during plating and staining of the screening library. Cells were pre-filtered according to their autophagy score using classifier version P where indicated.

| Classifier Version | Description | Architecture | Number of trainable parameters | Training data | Training Epochs | Number of cells per class | Fig. | Used to classify dataset | Independent Test Dataset | AUC ROC Curve |
| --- | --- | --- | --- | --- | --- | --- | --- | --- | --- | --- |
| **1.0** | Trained on initial staining protocol. Used to classify pilot screen | As in Fig. 2b but classifier head only consists of 3 fully connected linear layers | 17,882,244 | 2 slides unstimulated wt<br>2 slides wt + Torin-1<br>1 slide *ATG5*<sup>-/-</sup> clone 1<br>1 slide *ATG5*<sup>-/-</sup> clone 2 | 40 | 500,000 | 2, S2 | A<br>1.2 million cells | 1 slide *ATG5*<sup>-/-</sup> cells clone 1<br>1 slide unstimulated wt cells<br>1 slide wt cells + Torin-1 | 0.925879 |
| **P** | Only used to pre-filter cells for training 2.0 & 2.1 | As in first classifier | 17,882,244 | 3 slides wt + Torin-1<br>3 slides *ATG5*<sup>-/-</sup> mixed clones<br>1 slide *ATG5*<sup>-/-</sup> clone 1<br>1 slide *ATG5*<sup>-/-</sup> clone 2 | 30 | 1,200,000 |  |  | 1 slide *ATG5*<sup>-/-</sup> cells clone 1<br>1 slide unstimulated wt cells<br>1 slide wt cells + Torin-1 |  |
| **2.0** | Used on initial batch of genome-wide screen | As shown in Fig. 2b | 14,340,484 | 1 slide *ATG5*<sup>-/-</sup> clone 1<br>1 slide *ATG5*<sup>-/-</sup> clone 2<br>3 slides *ATG5*<sup>-/-</sup> mixed clones<br>2 slides unstimulated screen cells score > 0.9 (autophagy-off)<br>3 slides wt + Torin-1 score < 0.1 (autophagy-on) | 20 | 1,000,000 | 3, S4 | Screen batch 1<br>5 million cells | 1 slide *ATG5*<sup>-/-</sup> cells clone 1<br>1 slide unstimulated wt cells<br>1 slide wt cells + Torin-1 | 0.999649 |
| 2.1 | Refined for largest part of genome-wide screen | As shown in Fig. 2b | 14,340,484 | 1 slide *ATG5*<sup>-/-</sup> clone 1<br>1 slide *ATG5*<sup>-/-</sup> clone 2<br>3 slides *ATG5*<sup>-/-</sup> mixed clones<br>2 slides unstimulated screen cells score > 0.999 (autophagy-off)<br>3 slides wt + Torin-1 score < 0.001 (autophagy-on) | 20 | 1,000,000 | 4, S5 | Screen batch 2<br>35 million cells | 1 slide *ATG5*<sup>-/-</sup> cells clone 1<br>1 slide unstimulated wt cells<br>1 slide wt cells + Torin-1 | 0.999772 |

**Table S2.** Comparison of high-throughput methods for combined spatial phenotyping and genotyping Search space: library size that can be screened for phenotypes. Target space: Proportion of library that can be analyzed. Phenotypic variants that can be discriminated: The maximum number of different phenotypes that can be recovered from a single screen. Real time decision for genotyping necessary: Whether a decision has to be made for a given image in real time during screening (“yes”) or whether entire single cell datasets can be analyzed after imaging before a decision on which cells to genotype has to be made (“no”). *A genome-wide screen using in situ-seq has recently been reported (ref 21) with a small library of 10 million cells in which the number of screened and successfully sequenced cells and sgRNA representation remain unclear. °These technologies have low costs per screened cell, but require the use of instruments often provided by core facilities such as a laser dissection microscope for SPARCS, an imaging sorter device for imaging flow cytometry or an imaging setup equipped with a fluorescence recovery after photobleaching (FRAP) laser for pA-mCherry.

|  | SPARCS | In situ seq | Imaging flow cytometry | pA-mCherry |
| --- | --- | --- | --- | --- |
| References | This study | 19, 20, 21 | 22, 23 | 25, 26, 27 |
| Search space | large | medium | large | large |
| Target space | small | medium | large | small |
| Phenotypic variants that can be discriminated | microtiter plate | unlimited | microtiter plate | 4 per fluorophore |
| Image quality | confocal | confocal | flow-based | confocal |
| Real time decision for genotyping necessary | No | No | Yes | Yes |
| Discovery of new phenotypes after screen by reanalysis with new computational model possible | Yes | Yes | No | No |
| Largest library size | 40 million | 31 million | 12 million | 12.6 million |
| Genes targeted | 19,114 | 5,072* | 18,408 | 18,905 |
| Special equipment required | Laser microdissection microscope | Ultrafast imaging setup and precise stage for sequencing cycles | Imaging flow sorter | FRAP laser or equivalent |
| Cost per cell in library | Low° | High | Low° | Low° |

